# Functional genomics of a symbiotic community: shared traits in the olive fruit fly gut microbiota

**DOI:** 10.1101/590489

**Authors:** Frances Blow, Anastasia Gioti, Ian B. Goodhead, Maria Kalyva, Anastasia Kampouraki, John Vontas, Alistair C. Darby

## Abstract

The olive fruit fly *Bactrocera* oleae is a major pest of olives worldwide and houses a specialized gut microbiota dominated by the obligate symbiont “*Candidatus* Erwinia dacicola”. *Ca*. E. dacicola is thought to supplement dietary nitrogen to the host, with only indirect evidence for this hypothesis so far. Here, we sought to investigate the contribution of the symbiosis to insect fitness and explore the ecology of the insect gut. For this purpose, we examined the composition of bacterial communities associated with Cretan olive fruit fly populations, and inspected several genome and transcriptome assemblies. We identified, and reconstructed the genome of, a novel component of the gut microbiota, *Tatumella* sp. TA1, which is stably associated with Mediterranean olive fruit fly populations. We also reconstructed a number of pathways related to nitrogen assimilation and interaction with the host. The results show that, despite variation in taxa composition of the gut microbial community, core functions related to the symbiosis are maintained. Functional redundancy between different microbial taxa was observed for genes allowing urea hydrolysis. The latter is encoded in the obligate symbiont genome by a conserved urease operon, likely acquired by horizontal gene transfer, based on phylogenetic evidence. A potential underlying mechanism is the action of mobile elements, especially abundant in the *Ca*. E. dacicola genome. This finding, along with the identification, in the studied genomes, of extracellular surface structure components that may mediate interactions within the gut community, suggest that ongoing and past genetic exchanges between microbes may have shaped the symbiosis.

## Introduction

Many insects house gut microbial communities that perform essential functions related to their diet or lifestyle (Dillon and Dillon 2004). Insect species with a dependence on a specialized gut microbial community for their fitness often have specialized alimentary structures to house microbes, and these microbes are also often vertically transmitted between mother and offspring to ensure the inoculation of subsequent generations (Salem et al. 2015). The gut microbiota of wild populations of the olive fruit fly *Bactrocera oleae* (Tephritidae) is numerically dominated by a single Gammaproteobacterium, “*Candidatus* Erwinia dacicola” (Capuzzo et al. 2005; Estes et al. 2009; Kounatidis et al. 2009; Estes et al. 2012; Ben-Yosef et al. 2015). The symbiont *Ca.* E. dacicola resides in the digestive tract throughout the insect’s lifecycle, in larvae within the midgut caeca and in adults in a specialized foregut diverticulum structure called the esophageal bulb, and is maternally transmitted between generations by egg smearing (Estes et al. 2009). *Ca.* E. dacicola has not been detected outside of the insect by environmental sampling, while efforts to culture it *in vitro* have so far been unsuccessful (Capuzzo et al. 2005). From the above, the bacterium qualifies as a specific obligate symbiont of *B. oleae,* but also a challenge to study experimentally. In this context, sequence data-based methods are of high value for the generation of hypotheses on the different roles of *Ca.* E. dacicola in host fitness.

In common with many holometabolous insects, the adult and juvenile stages of *B. oleae* feed on different food sources (Tzanakakis 2003), therefore the *Ca.* E. dacicola symbiosis is hypothesized to perform alternative functions during these developmental stages. In larvae, the symbiosis has been hypothesized to allow the insect to use ripening fruit by detoxifying defensive plant phenolic compounds, such as oleuropein (Soler-Rivas et al. 2000; Ben-Yosef et al. 2015), and sequestering amino acids for protein synthesis (Ben-Yosef et al. 2015). During adulthood, the native gut microbiota is thought to provision nitrogen to the host: females harboring their native microbiota have a higher reproductive output on diets lacking essential amino acids (EAAs), and diets containing only nitrogen sources that are metabolically intractable to *B. oleae,* such as urea (Ben-Yosef et al. 2010; Ben-Yosef et al. 2014). It is common for many insects to house microbes that increase the quantity or quality of dietary nitrogen by performing novel metabolic functions (Douglas 2009). For example, the intracellular symbiont *Blochmannia* within the bacteriocytes of *Camponotus* ants encodes urease that metabolizes dietary urea to ammonia from which it synthesizes essential and non-essential amino acids, subsequently transported to the hemolymph for host consumption (Feldhaar et al. 2007). A similar process is proposed to occur in *B. oleae* adults, which consume bird droppings containing ammonia and urea as part of their omnivorous diet (Ben-Yosef et al. 2014), but do not encode endogenous ureases (*B. oleae* genome accession number: GCF_001188975.1, Djambazian et al. 2018). However, the specific pathways and related enzymes involved in the metabolism and uptake of nitrogenous substrates remain to be elucidated in *B. oleae.*

Previous experimental approaches aiming to evaluate symbiotic function have tested the native microbial community of the olive fruit fly, which is dominated by, but not restricted to *Ca.* E. dacicola (Estes et al. 2009; Ben-Yosef et al. 2015; Blow, Vontas, et al. 2016; Blow et al. 2017). In this study, we took advantage of recently available –omic resources for the symbiont (Blow, Gioti et al. 2016, Pavlidi et al. 2017, Estes et al, 2018) and further generated new sequence data from single-culture and metagenomic samples of the *B. oleae* gut community to 1) investigate whether *Ca.* E. dacicola encodes the full gene repertoire required to provide dietary nitrogen to *B. oleae* and 2) identify other members of the *B. oleae* gut microbiota potentially capable of performing these functions. For this purpose, we employed comparative genomic and phylogenetic analyses, considering the function of the gut community as a whole in the context of symbiosis.

## Results and discussion

### A novel member of the gut microbiota associated with Mediterranean populations of *B. oleae*

To explore the ecology of the *B. oleae* gut, we examined its bacterial community composition by 16S rRNA gene amplicon sequencing of samples from olive fruit fly populations collected in Crete. This approach, combined with culturing of selected isolates, allowed identifying and isolating in to axenic culture a culture-viable member of the gut microbiota that has not been characterized before, referred to here as *Tatumella* sp. TA1. *Tatumella* sp. TA1 was initially identified in wild Cretan populations of adult *B. oleae,* and reanalysis of 16S rRNA gene data from a previous study demonstrated that *Tatumella* sp. TA1 is also present in wild populations from Israel (Ben-Yosef et al. 2015). Notably, *Tatumella* sp. TA1 was not detected in any of the studies of bacterial communities associated with US populations of *B. oleae* (Estes et al. 2009; Estes et al. 2012). These results suggest that TA1 is a facultative member of the gut community and that it is restricted to Mediterranean populations of the olive fruit fly. We detected *Tatumella* sp. TA1 in 88.9-90 % of individuals from populations in Israel, in both adults and larvae, and in 26.9 % of individuals from populations in Crete (Table 1). We did not detect *Enterobacter* sp. OLF, another facultative member of the gut community identified in US populations of *B. oleae* (Estes et al. 2009), in either of the Mediterranean populations studied, suggesting that the two taxa are geographically restricted.

**Table 1.**
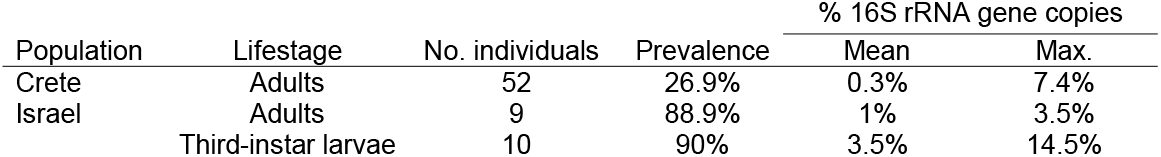
Prevalence of *Tatumella* sp. TA1 in Mediterranean populations of *B. oleae.*

We used PacBio RS II sequencing to generate a draft genome sequence for *Tatumella* sp. TA1. The assembly comprised a chromosomal sequence of 3,389,139 bp and a circular plasmid sequence of 49,211 bp (total genome size 3.4 Mb) encoding a total of 3,309 protein coding genes, 72 tRNAs, and 22 rRNAs. Both the chromosome and plasmid sequences have been circularized. The GC content of the genome was 48.6 %, and both this and the genome size were comparable to those of previously-published genomes of *Tatumella* species, many of which are host-associated (Hollis et al. 1981; Marín-Cevada et al. 2010; Chandler et al. 2014). The assembly is predicted to be 100 % complete following the method used in (Rinke et al. 2013), which detected 138/138 single-copy marker genes.

### Phylogenetic placement of obligate and facultative members of the *B. oleae* gut microbiota

In agreement with a previously-published comprehensive phylogeny (Palmer et al. 2017), ortholog analysis of 46 bacterial taxa belonging to the genera *Erwinia, Tatumella* and *Pantoea* (full list in Table S1A) supports the phylogenetic placement of *Tatumella* sp. TA1 within the *Tatumella* genus (Figure 1). Phylogenetic analyses confirmed that *Tatumella* sp. TA1 and *Enterobacter* sp. OLF are distinct members of the order *Enterobacterales,* indicating that they are unique components of the *B. oleae* gut microbiota. In addition, the phylogeny showed that, despite an evolutionary association with *B. oleae* and vertical transmission, which are expected to change genome characteristics through altered selection pressures and drift (McCutcheon and Moran 2011), *Ca.* E. dacicola clusters within the *Erwinia* genus (Figure 1). The closest phylogenetic relatives of *Tatumella* sp. TA1 and *Ca.* E. dacicola are the thrip symbionts BFo2 and BFo1, respectively (Facey et al. 2015). The clustering of these taxa may represent comparable lifestyles. In a result analogous to diet experiments in olive flies (Ben-Yosef et al. 2010; Ben-Yosef et al. 2014), the fitness benefits of the WFT gut-lumen symbiont BFo1 were condition-dependent and only apparent on diets lacking a balanced source of amino acids, when gut bacteria are presumed to synthesize and provision adult WFT with amino acids required for protein synthesis (de Vries et al. 2004). BFo1 and BFo2 also inhabit the gut lumen and are vertically transmitted between generations by an extracellular route (de Vries et al. 2001). Further functional studies may elucidate whether specific characteristics shared by these taxa predispose them to this lifestyle.

**Fig. 1.**
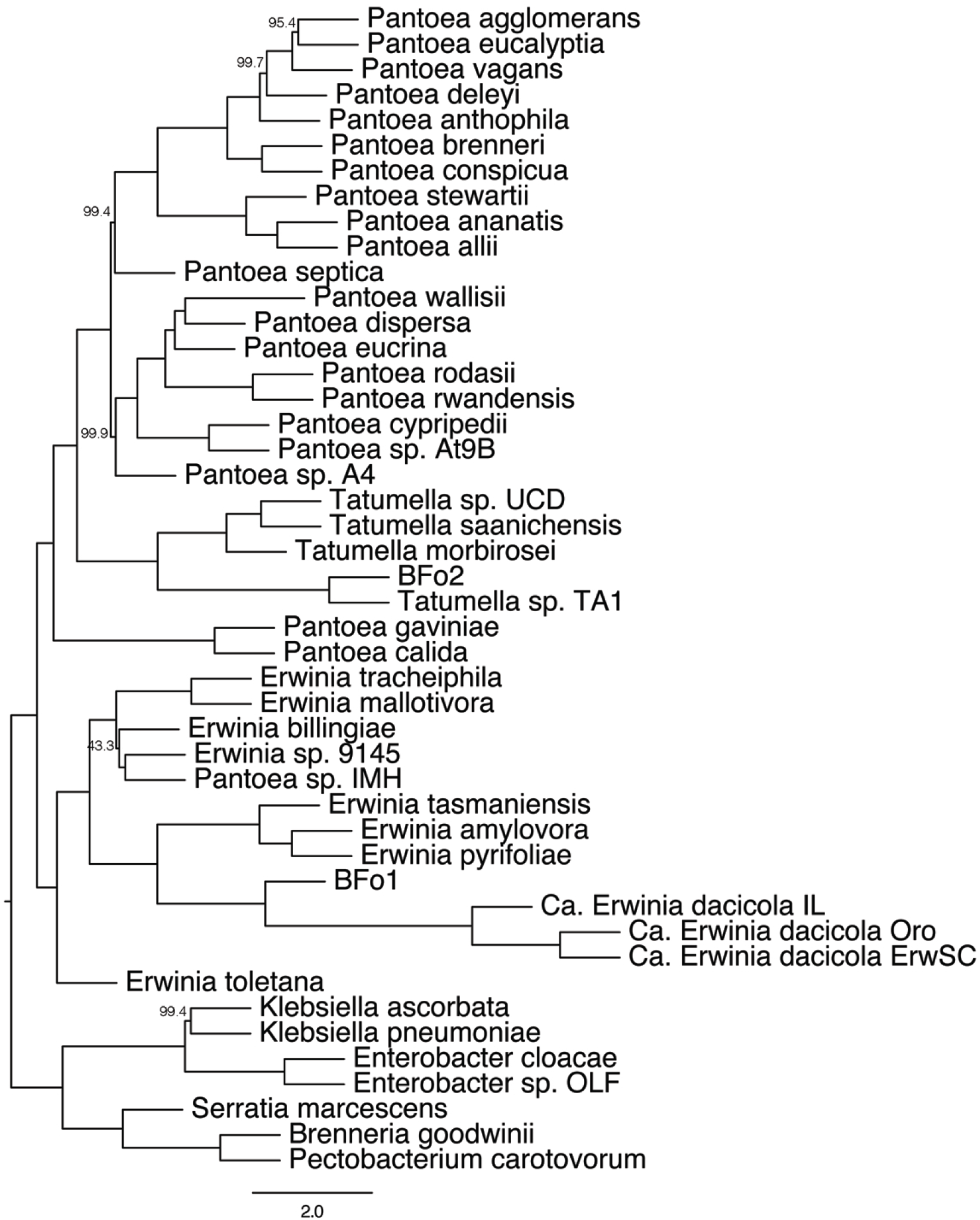
Maximum-Likelihood tree estimated from amino acid alignments of 289 single-copy orthologs retrieved from whole genome assemblies of 39 *Erwinia, Pantoea* and *Tatumella* and 7 outgroup taxa. Full names and accession numbers of the genes used are available in Table S1A. Numbers on nodes correspond to bootstrap support (only values < 100 are shown).

### Functional redundancy of nitrogen assimilation genes in members of the *B. oleae* gut microbiota

We investigated the hypothesis that members of the *B. oleae* gut microbiota supplement the olive fruit fly diet with nitrogen by focusing on both obligate and facultative bacterial members. The annotated genomes of the obligate *Ca.* E. dacicola (Blow, Gioti et al. 2016) and facultative *Tatumella* sp. TA1 (this study) and *Enterobacter* sp. OLF (Estes et al. 2018) all encode enzymes that metabolize urea to ammonia and carbon dioxide (Figure 2, Table S2A): *Ca.* E. dacicola and *Enterobacter* sp. OLF encode ureases (EC 3.5.1.5), and *Tatumella* sp. TA1 encodes urea carboxylase (EC 6.3.4.6) and allophanate hydrolase (EC 3.5.1.54). None of these enzymes are encoded in the genome of *B. oleae* (Table S2A). A BlastP and TblastN search of the *Ca.* E. dacicola transcriptome data from larvae developing in green and black olives (Pavlidi et al. 2017) did not identify any of the urease genes, indicating that urease may not be expressed at the larval stage in the obligate symbiont. This result is expected, since at this stage there is no dietary source of urea. Therefore, a hypothesis to further explore is that urease expression in *Ca.* E. dacicola is inducible by host or dietary cues present during *B. oleae* adulthood. Similarly, the closest homologs to the *Ca.* E. dacicola urease gene (see next section) come from two plant pathogens, *Gibbsiella quercinecans* and *Brenneria goodwinii,* previously characterized as negative for urease function (Brady et al. 2010; Denman et al. 2012), which indicates that urease function may be inducible under specific conditions.

**Fig. 2.**
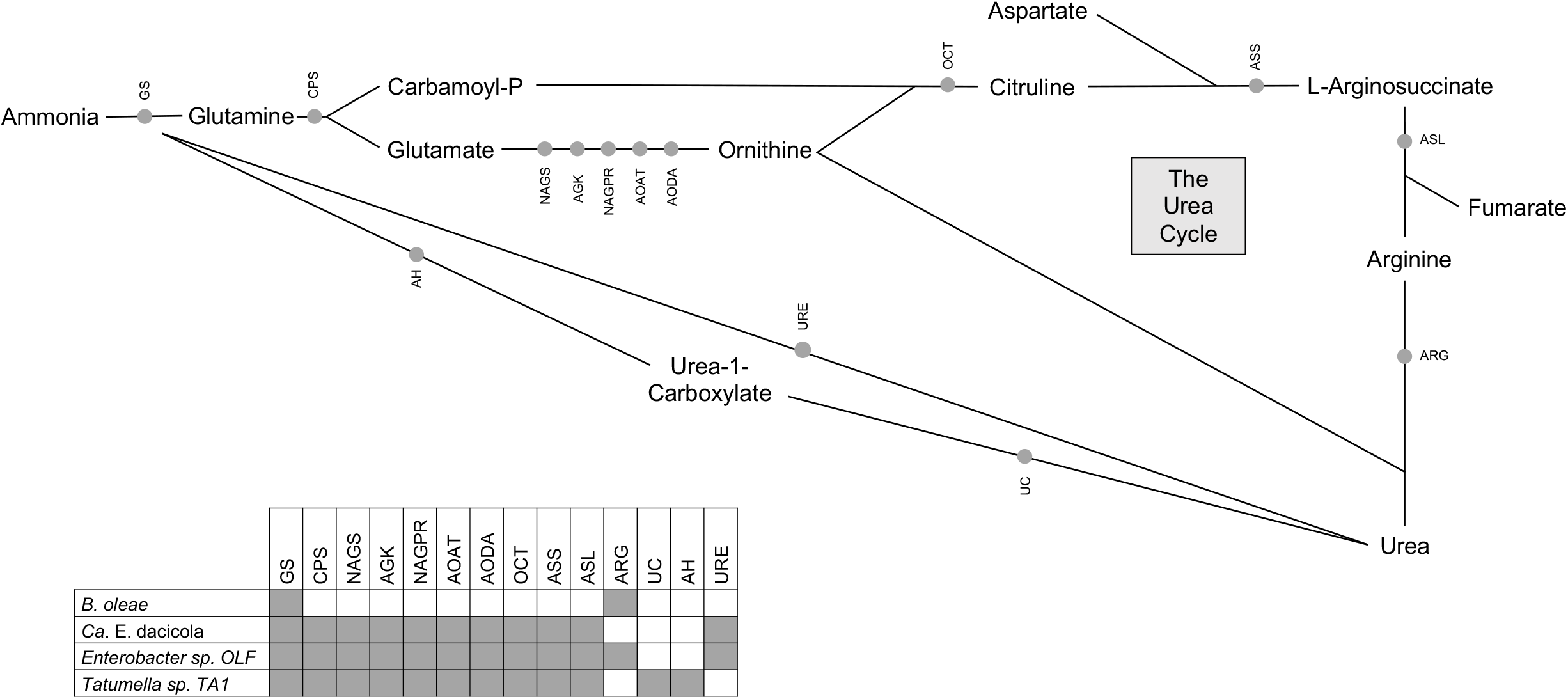
Proposed pathways for urea hydrolysis and recycling of nitrogen into host and microbial metabolic pathways via ammonia, glutamine and glutamate. The genes encoding the illustrated enzymes were detected in genome assemblies of *Ca.* E. dacicola (Blow, Gioti, et al. 2016), *Tatumella* sp. TA1 (this study), both annotated with PROKKA, and *Enterobacter* sp. OLF annotated as described in (Estes, Hearn, Nadendla, et al. 2018). GS (Glutamine synthetase E.C. 6.3.1.2); CPS (Carbamoyl-phosphate synthase E.C. 6.3.4.16; E.C. 6.3.5.5); NAGS (N-acetylglutamate synthase E.C. 2.3.1.1); AGK (Acetylglutamate kinase E.C. 2.7.2.8); NAGPR (N-acetyl-gamma-glutamyl-phosphate reductase E.C. 1.2.1.38); AOAT (Acetylornithine/succinyldiaminopimelate aminotransferase E.C. 2.6.1.11); AODA (Acetylornithine deacetylase E.C. 3.5.1.16); OCT (Ornithine carbamoyltransferase E.C. 2.1.3.3); ASS (Argininosuccinate synthase E.C. 6.3.4.5); ASL (Argininosuccinate lyase E.C. 4.3.2.1); ARG (Arginase E.C. 3.5.3.1); UC (Urea carboxylase E.C. 6.3.4.6); AH (Allophanate hydrolase E.C. 3.5.1.54); URE (Urease E.C. 3.5.1.5).

In addition to urea hydrolysis, *Ca.* E. dacicola, *Tatumella* sp. TA1 and *Enterobacter* sp. OLF all encode biosynthetic operons for non-essential and essential amino acids (EAA), with further evidence for expression during larval feeding in *Ca.* E. dacicola (Table S2B, sequences available in Table S3). Glutamine synthetase (EC 6.3.1.2), is also present in all three taxa and expressed in the *Ca.* E. dacicola transcriptome data. This enzyme, also encoded by the host genome (RefSeq accession GCF_001188975.1), incorporates nitrogen from ammonia into the non-essential amino acid glutamine (Figure 2), which can be channeled into downstream metabolic processes including EAA biosynthesis (Feldhaar et al. 2007; Sabree et al. 2009).

We also investigated the hypothesis that host nitrogenous waste, which would otherwise be excreted, is the source of nitrogen assimilated to non-essential amino acids by the gut microbiota. Urea could come from recycling of host waste uric acid (Ben-Yosef et al. 2014), as has been observed in other insect symbionts that upgrade the nitrogen content of the host diet (Potrikus and Breznak 1981; Sasaki et al. 1996; Kashima et al. 2006; Feldhaar et al. 2007; Sabree et al. 2009). We found no genomic evidence that urea can be produced via uricolysis in either the host or any member of the so-far identified and sequenced microbiota: BlastP searches against shotgun sequencing data of olive fruit fly gut samples and corresponding metagenomic assemblies of the gut microbial community (NCBI BioProject accession PRJNA326914) failed to identify the required enzyme allantoicase (EC 3.5.3.4; Figure S1, Table S2A). However, *B. oleae* does encode an arginase (EC 3.5.3.1), which can hydrolyze the essential amino acid arginine to urea and ornithine and could provide urease-degrading bacteria with a supply of urea (Figure 2). One hypothesis would be that *B. oleae* might enrich its gut microbiota with ureolytic bacteria such as *Ca.* E. dacicola through inducible delivery of urea produced by arginase activity. Signal peptides were not detected when individual urease proteins or the full operon from *Ca.* E. dacicola were analyzed with SignalP version 4.0 (Petersen et al. 2011), indicating that, if expressed, the urease encoded by *Ca.* E. dacicola is intracellular. This suggests that specific uptake of urea and excretion of the products of its hydrolysis must either be coordinated with the host, or that microbial cells are lysed after the hydrolysis of urea, in order for the host to gain the observed nutritional benefit when urea is added to the diet (Ben-Yosef et al. 2010; 2014). A key component of future studies should be to track the fate of metabolites resulting from urea hydrolysis: whether they are retained by microbes for endogenous metabolism, or whether they are trafficked back to the host as e.g. ammonia or glutamine.

### Evidence for horizontal gene transfer of the urease operon in the obligate symbiont *Ca.* E. dacicola

The *Ca.* E. dacicola urease enzyme is encoded, as in all ureolytic bacteria, by a cluster of eight adjacent genes arranged in an operon and denoted as *ureDABCEJFG,* with *ureABC* encoding structural proteins of the enzymatic complex, *ureDEFG* accessory proteins, and *ureJ* the transcriptional regulator. The urease operon appears incomplete at its 3’ end in the ErwSC assembly (Blow, Gioti et al. 2016), missing *ureG.* However, *ureG* is present in the *Ca.* E. dacicola genome, as confirmed by BlastP queries against the initial set of metagenomic reads obtained from shotgun sequencing (NCBI BioProject accession PRJNA326914). The urease operon is also present in two additional *Ca.* E. dacicola assemblies (IL and Oroville) reconstructed with different methods and from different olive fly populations (Table S1A), and is complete (5,962 bp total length) in these assemblies, including *ureG.* The three ‘versions’ of the operon from different assemblies are 99% similar at the nucleotide level for the genes present. One explanation for the absence of *ureG* in the ErwSC assembly is over-trimming, aiming to minimize the risk of including non-Ca. E. dacicola scaffolds, since metagenomic sequencing data included DNA from several species, such as *Tatumella* sp. TA1. The differences between the reduced ErwSC and the more complete Oroville assembly are outlined in detail in (Estes, Hearn, Agrawal, et al. 2018). In any case, the presence and similarity of the operon in all three *Ca* E. dacicola assemblies despite their differences and despite the presence of different facultative taxa in the gut microbiome of *B. oleae* from distinct geographic populations, indicate that the urease operon is highly conserved in *Ca* E. dacicola.

To the best of our knowledge, *Ca.* E. dacicola is the only species of the genus *Erwinia* that encodes urease genes. Examination of the phylogeny of *ureC,* the longest structural gene of the operon, provided evidence for the acquisition of the urease operon in *Ca.* E. dacicola by horizontal gene transfer (HGT): The *Ca.* E. dacicola *ureC* history (Figure 3A), is different from that of the species history, as shown by the comparison to a ‘reference’ tree, reconstructed from 8 neutral single-copy orthologs (Figure 3B). The reference tree accurately depicts the known phylogenetic relationships between *Ca.* E. dacicola and other Proteobacteria, in agreement with the recently published comprehensive phylogeny (Palmer et al. 2017) and the species tree presented in this study (Figure 1). By contrast, the *Ca.* E. dacicola *ureC* protein groups together with proteins from more distant Gammaproteobacteria of the *Enterobacterales* order (*G. quercinecans, B. goodwinii*), and members of the distantly related Betaproteobacteria (Figure 3). Hacker and Kaper (2000) defined genomic islands (GEIs) in bacteria as small (<10 Kbp), syntenic blocks of genes acquired by HGT. It has been observed that GEIs share some common characteristics, such as the encoding of genes offering a selective advantage to the host, flanking by mobile genetic elements (MGEs) such as transposases, insertion close to tRNA genes, and a GC content different from that of the rest of the genome (Juhas et al. 2009). The ureolytic capacity conferred to *Ca.* E. dacicola by the urease operon is potentially an example of acquired selective advantage, since it increases the availability of nutrients to the symbiont itself or to the host, upon which it is dependent. Thus, it is tempting to argue that the urease operon represents a GEI in *Ca.* E. dacicola. Similarly to other GEIs, it has a distinct (lower) GC content compared to the rest of the genome (mean GC_urease_=0.502, mean GC_genome_=0.535), and this difference is statistically significant (p = 5.085e-06). Moreover, the distribution of GC values sampled from the urease operon is significantly different from the whole-genome distribution (p = 1.031E-05). Unfortunately, proximity of the operon to MGEs and tRNAs could not be confirmed in *Ca.* E. dacicola, since its genomic architecture remains unresolved: The operon appears in a separate scaffold in all three available assemblies (e.g. IL assembly accession number: LJAM02000124.1), none of which have been further extended and ordered with the use of long insert-size libraries. MGEs are frequently associated with fragmented assemblies, and in the ErwSC assembly, we detected a transposase fragment upstream of the first gene of the operon, *ureD,* in the scaffold containing the urease. However, mate-pair library sequencing data from Cretan olive fruit fly guts (accession number: SRX1896451) did not support the presence of the transposase upstream of *ureD,* and did not allow identification of the operon’s flanking scaffolds.

**Fig. 3.**
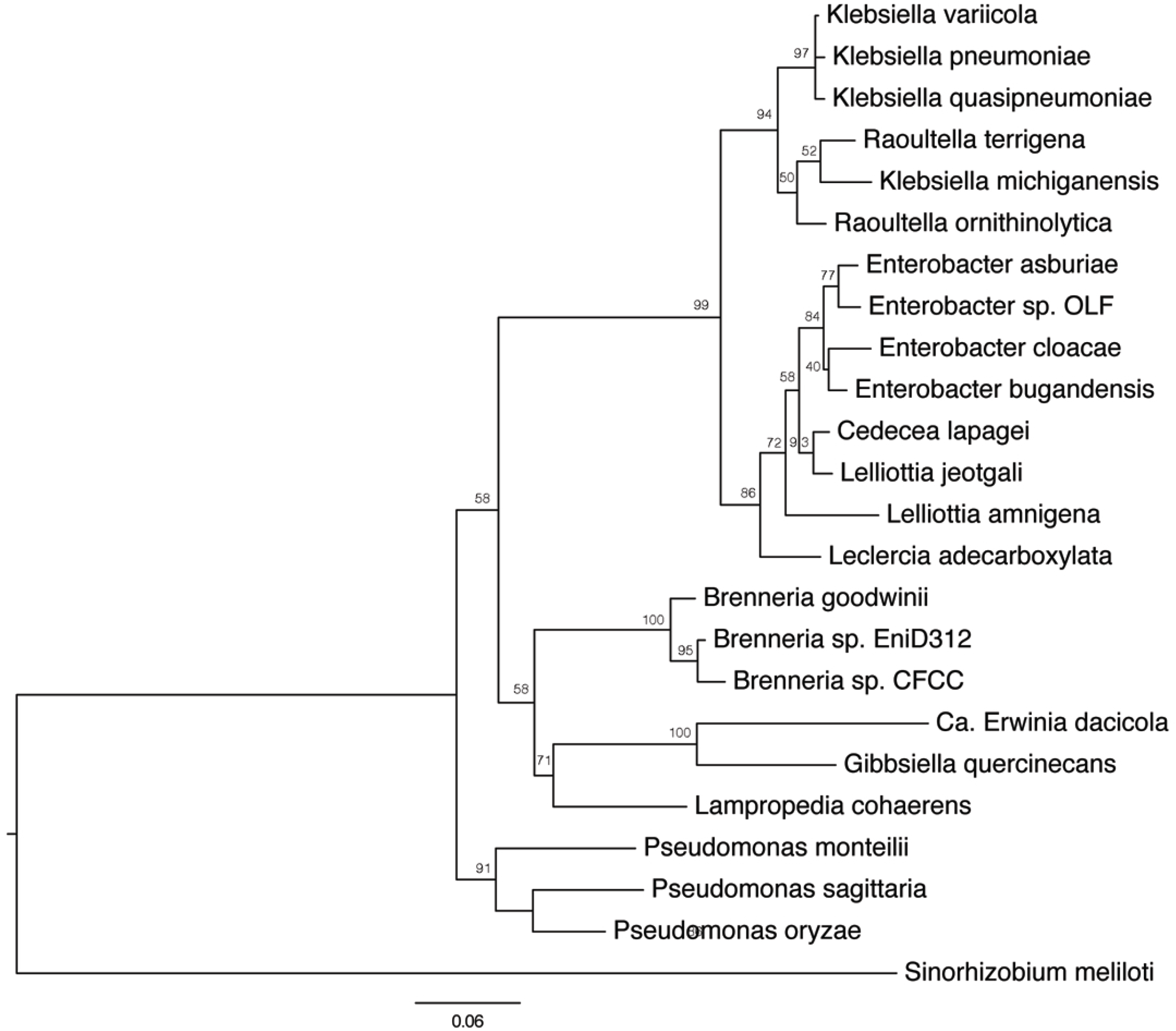
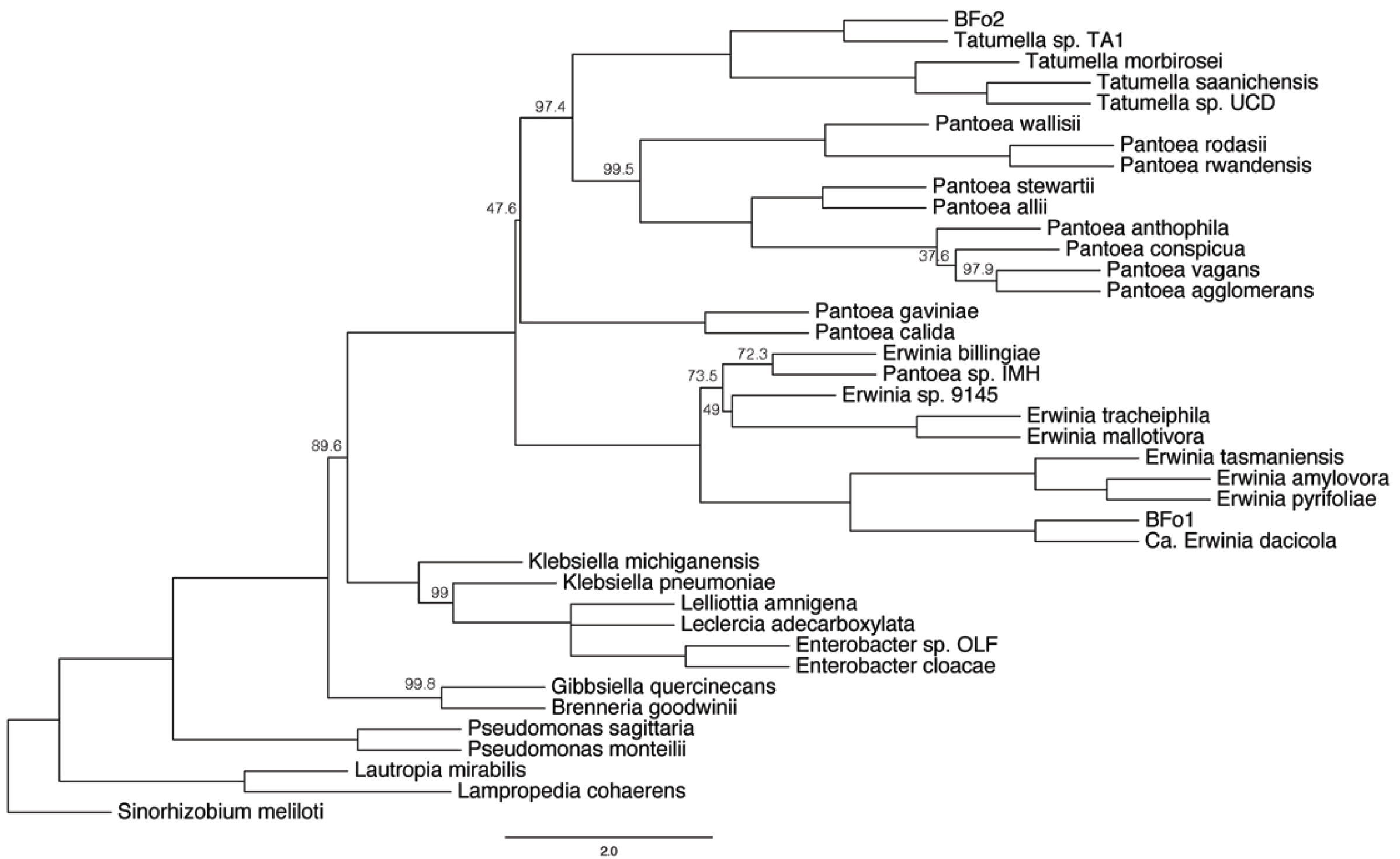
Maximum-Likelihood trees estimated from amino acid alignments of (*A*) the urease subunit alpha (*ureC*) gene and (*B*) 8 single-copy orthologs, representing a ‘neutral reference’. Rooting of both trees was performed with the most phylogenetically remote taxon, the Alphaproteobacterium *Sinorhizobium meliloti.* Full names and accession numbers for the genes (*A*) and genomes (*B*) used are available in Table S1B and Table S1A, respectively. Numbers on nodes correspond to bootstrap support.

The present data do not allow us to identify the date of the acquisition of the urease operon, nor its taxonomic origin. The observation that the GC content of the operon is distinct from that of its host argues in favor of a relatively recent acquisition, as previously proposed for a different *Ca.* E. dacicola GEI (Estes et al. 2018), but one cannot conclude without dating the symbiosis. Similarly, regarding the donor, the low bootstrap support for the clades that link the groups comprising *Ca.* E. dacicola, *G. quercinecans* and *Brenneria* to the rest of the clades indicates that the original donor of the urease cluster might be extinct or not yet sequenced. One possibility is that it is a free-living bacterium encountered by *Ca.* E. dacicola before its transition to obligacy with *B. oleae* or a yet-uncharacterized gut bacterium. In line with the latter, there is abundant evidence that members of gut microbial communities in vertebrates and invertebrates undergo HGT, and that this can subsequently influence functional traits (Petridis et al. 2006; Smillie et al. 2011; Stecher et al. 2012; Baker et al. 2018). Linked to HGT events, mobile genetic elements (MGEs), including transposases, repetitive DNA regions and phages, are abundant in all three *Ca.* E. dacicola assemblies: MGEs represent an average 34 % of the obligate symbiont’s genome, in contrast to the corresponding MGE content in *Tatumella* sp. TA1 and *Enterobacter* sp. OLF (Table S4). However, these percentages should not to be taken at absolute value due to the inherent difficulties in computational annotation of MGEs, especially from metagenomic samples (Jørgensen et al. 2014), while they might represent an overestimate due to the draft - and often fragmented - nature of the assemblies. Still, it is notable that high transposase content has been often observed in bacteria that recently adopted a symbiotic lifestyle (Gil et al. 2008; Ran et al. 2010), while MGEs were previously shown to be transcriptionally active in *B. oleae* larvae feeding on green and black olives (Pavlidi et al. 2017). These findings overall support the idea that genetic exchange between *Ca.* E. dacicola and members of the *B. oleae* gut microbial community may occur.

### Genes encoding extracellular surface structure components in the *B. oleae* gut microbiota

Exchanges of genetic material between bacteria are facilitated by extracellular surface structures (Thomas and Nielsen 2005), components of which are encoded in the genomes of the *B. oleae* gut microbiota: Components of the IncF transfer system of conjugative plasmids, encoding F-like pili, are present in the two most complete *Ca.* E. dacicola assemblies, IL and Oroville (Table S5). In addition, the chromosome of *Tatumella* sp. TA1 encodes a Trb operon for conjugative transfer, potentiating the exchange of genetic material (Table S5). Notably, plasmid pTA1 from *Tatumella* sp. TA1, identified in the present study (GenBank accession number: CP033728), is present in the *Ca.* E. dacicola Oroville assembly, which was generated from *B. oleae* populations where *Tatumella* sp. TA1 has not yet been detected. This result, most likely not due to ‘contamination’ as confirmed by the absence of 16S rRNA gene sequences from taxa other than *Ca.* E. dacicola in the Oroville assembly (data not shown), indicates that transfers of genetic material among members of the *B. oleae* gut microbiota occur.

Some extracellular surface structures that mediate the exchange of genetic material between bacteria, such as type IV secretion systems (T4SS), also facilitate adhesion and molecular exchange with eukaryotic cells (Alvarez-Martinez and Christie 2009). T4SS are diverse; VirB-like T4SS encode short and rigid pili, whereas the F-pili T4SS for conjugative transfer tend to be long and flexible (Clarke et al. 2008). Depending on their structure, pili can deliver a range of effector molecules including proteins and nucleic acids to both prokaryotic and eukaryotic cells, and can also be involved in the transition between motility and sessility, for example in the process of biofilm formation (Christie and Vogel 2000). T4SS mediate the delivery of effector proteins directly to the cytoplasm of eukaryotic cells during the process of infection in pathogenic bacteria (Cascales and Christie 2003), and root nodule colonization in symbiotic *Rhizobia* (Deakin and Broughton 2009). Effector proteins can induce regulatory changes in the eukaryotic cell, triggering modification of cellular conditions or conditions in the environment to facilitate growth or invasion (Burns 2003). A conserved VirB-like T4SS operon encoding the essential genes for pilus formation and substrate trafficking (VirB1-11 and VirD4, (Wallden et al. 2010)) was detected in all three *Ca.* E. dacicola assemblies, but was not detected in either *Enterobacter* sp. OLF or *Tatumella* sp. TA1 (Table S5). Evidence for expression of the *Ca.* E. dacicola VirB-like T4SS operon during *B. oleae* juvenile development in ripening olives (Pavlidi et al. 2017) and for the presence of pili-like structures in the esophageal bulb (Poinar et al. 1975) suggest that the T4SS system pili may also play a - yet unknown- role in the *B. oleae* - *Ca.* E. dacicola symbiosis, for example by the establishment of *Ca.* E. dacicola biofilms (Estes et al. 2009). One potential function to explore in future studies might be the coordination of gut lumen colonization and subsequent resource exchange between the host and symbiont.

## Conclusions

In this study, we aimed to gain understanding on the advantages of symbiosis for the olive fruit fly *B. oleae* by integrating various sources of genetic information on its microbial gut community. This led to the identification, by 16S rRNA gene sequencing and culture-dependent methods, of a novel facultative member of the gut microbiota, *Tatumella* sp. TA1, which exclusively associates with Mediterranean populations of *B. oleae* throughout the lifecycle. *Tatumella* sp. TA1 is phylogenetically distinct from the US-restricted facultative symbiont *Enterobacter* sp. OLF, highlighting the population-dependent nature of gut microbiota composition in the olive fruit fly. Comparative genomics indicated that the obligate symbiont *Ca.* E. dacicola, as well as *Tatumella* sp. TA1 and *Enterobacter* sp. OLF, which are all stable components of the *B. oleae* microbiota throughout the olive fruit fly lifecycle, encode genes that allow the use of urea as a nitrogen source. The hydrolysis of urea to ammonia is encoded by genes with distinct phylogenetic origins in each organism: HGT of a urease operon in *Ca.* E. dacicola, an endogenous urease operon in *Enterobacter* sp. OLF, and the presence of alternative enzymatic machinery (urea carboxylase and allophanate hydrolase) in *Tatumella* sp. TA1. These findings provide a potential mechanistic basis for previous experimental evidence of gut microbiota-mediated dietary nitrogen provisioning during adulthood, but emphasize the need for experimental validation of metabolic cross-feeding between the host and obligate symbiont. A hypothesis in this direction, warranting focus in future studies, is that a VirB-like T4SS encoded by *Ca.* E. dacicola, but missing from the genomes of *Enterobacter* sp. OLF and *Tatumella* sp. TA1, may facilitate specific interactions with the host at the symbiotic interface. Besides T4SS, our study further highlighted extracellular surface structures, encoded in the genomes of the obligate and the facultative symbionts, as mediators of DNA transfer: Detection of a plasmid shared by geographically-distinct *Ca.* E. dacicola and *Tatumella* sp. TA1, along with the horizontal acquisition of genes important to symbiosis function (urease in this study, genes related to amino-acid degradation in (Estes, Hearn, Agrawal, et al. 2018)) and the abundance of mobile genetic elements in the obligate symbiont genome, indicate that previous and on-going genetic exchanges between gut community members are important determinants of symbiotic interactions with *B. oleae.*

## Materials and Methods

### Insect material

*B. oleae* adults were obtained by collecting infested olives from trees in the grounds of the University of Crete (Heraklion, Greece) in October and November of 2014. Infested olives were suspended over sterile sand with a wire mesh, and third-instar larvae were allowed to emerge from the fruit. Flies pupated in the sand and were placed into Petri dishes in 10 cm^3^ plastic cages prior to emergence. Flies were maintained at 25 °C and 60 % relative humidity and were supplied with artificial diet (19 % hydrolyzed yeast, 75 % icing sugar, 6 % egg yolk). Each cage was provided with Milli-Q water in a clean plastic container and wax cones for oviposition.

### Preparation and sequencing of 16S rRNA gene amplicon libraries

DNA was extracted from a total of 52 whole *B. oleae* adults using the Qiagen DNeasy Blood and Tissue kit for Gram-positive bacteria (Qiagen, UK) following the manufacturer’s instructions. An additional bead-beating step using 3 mm carbide beads (Qiagen, UK) in a Qiagen tissue lyzer (Qiagen, UK) at 25 Hz for 30 seconds was employed. In order to assess the diversity and composition of the bacterial communities, bacterial 16S rRNA V4 regions were amplified by PCR using the universal primers F515 (5’-GTGCCAGCMGCCGCGGTAA-3’) and R806 (5’- GGACTACHVGGGTWTCTAAT-3’) (Caporaso et al. 2011). Samples were dual-indexed for sequencing following the method in D’Amore et al. (2016). PCR reactions were performed in a total volume of 20 μl, containing 5 ng of template DNA, 10 μl NEBNext 2x High-Fidelity Master Mix (New England Biolabs), 0.3 μM of each primer, and 3.4 μl PCR-clean water. Thermal cycling conditions were 98 °C for 2 minutes, 10 cycles of 98 °C for 20 seconds, 60 °C for 15 seconds, and 70 °C for 30 seconds, with a final extension at 72 °C for 5 minutes. PCR products were purified with Agencourt AMPure XP beads (Beckman Coulter Genomics) and used as template for the second PCR reaction. Purified first-round PCR products were combined with 10 μl NEBNext 2x High-Fidelity Master Mix (New England Biolabs), and 0.3 μM of each barcoding primer containing adapters and indexes to a total volume of 20 μl. Thermal cycling conditions were 98 °C for 2 minutes, 15 cycles of 98 °C for 20 seconds, 55 °C for 15 seconds and 72 °C for 40 seconds, with a final extension at 72 °C for 60 seconds. PCR products were purified with Agencourt AMPure XP beads (Beckman Coulter Genomics) and quantified with the Qubit dsDNA High-Sensitivity assay (Life Technologies), and an Agilent Bioanalyzer High-Sensitivity DNA chip (Agilent). Samples were pooled at equimolar concentrations and size-selected in a range of 350-450 bp by Pippin-Prep (Sage Science). Sequencing was performed at the University of Liverpool Centre for Genomic Research on an Illumina MiSeq platform with V2 chemistry, generating 250 bp paired-end reads. All raw sequencing reads were deposited at NCBI under the BioProject accession PRJNA321174.

### Computational analyses of 16S rRNA sequencing data

Raw sequencing reads were de-multiplexed and converted to FASTQ format using CASAVA version 1.8 (Illumina 2011). Cutadapt version 1.2.1 (Martin 2011) was used to trim Illumina adapter sequences from FASTQ files. Reads were trimmed if 3 bp or more of the 3’ end of a read matched the adapter sequence. Sickle version 1.200 (Joshi & Fass 2011) was used to trim reads based on quality: any reads with a window quality score of less than 20, or were less than 10 bp long after trimming, were discarded. BayesHammer was used to correct reads based on quality (Nikolenko et al. 2013). Paired-end reads were merged with a minimum overlap of 50 bp using PandaSeq (Masella et al. 2012). All subsequent analyses were conducted in QIIME version 1.8.0 (Caporaso, Kuczynski, et al. 2010). Sequences were clustered into Operational Taxonomic Units (OTUs) by *de novo* OTU picking with USEARCH (Edgar 2010). Chimeras were detected and omitted with UCHIME (Edgar et al. 2011) and the QIIME-compatible version of the SILVA 111 release database (Quast et al. 2013). The most abundant sequence was chosen as the representative for each OTU, and taxonomy was assigned to representative sequences by BLAST (Altschul et al. 1990) against the SILVA 111 release database, which was supplemented with several reference sequences for *Ca.* E. dacicola. OTUs were filtered from the dataset if they matched the SILVA 111 database for chloroplasts or mitochondria. OTU representative sequences were aligned against the Greengenes core reference alignment (DeSantis et al. 2006) using PyNAST (Caporaso, Bittinger, et al. 2010). Previously published 16S rRNA gene amplicons from flies collected in Israel (Ben-Yosef et al. 2015) were employed for comparison with the above data. Since data generation methods for the samples from Israel varied slightly, all data analysis methods were standardized from read-error correction onwards in this study. *Tatumella* sp. TA1 prevalence was calculated as the proportion of individuals where *Tatumella* sp. TA1 16S rRNA was > 0.01% of the total 16S rRNA gene copies from that sample, and relative abundance was the proportion of the total 16S rRNA gene copies.

### Isolation, culture and identification of *Tatumella* sp. TA1 from *B. oleae*

*Tatumella* sp. TA1 was identified from a two-day old male fly following the below isolation and culture procedures, all conducted under sterile conditions using aseptic technique or in a laminar flow cabinet. Individual two-day old adult *B. oleae* collected in Crete were surface-sterilized with 70 % ethanol and rinsed twice in distilled water prior to homogenization with a plastic pestle in 20 μl nuclease-free water. 10 μl of the homogenate was spotted on to Columbia agar (Oxoid, UK) supplemented with 5 % defibrinated horse blood (TCS Biosciences, UK), and plates were incubated at 25 °C for 48 h. Single colonies were picked and streaked onto Brain Heart Infusion (BHI) agar (Oxoid, UK) to establish pure cultures. Pure liquid cultures were generated by inoculating single colonies from BHI agar plates in to BHI liquid medium and incubating them at 25 °C for 48 h. Liquid cultures were cryopreserved following the addition of a cryoprotectant (20 % [v/v] glycerol final concentration) and storage at −80 °C. In order to identify isolates, single colonies from BHI agar plates were picked and inoculated into 10 μl nuclease-free water in a PCR tube. Tubes were incubated at 95 °C for 5 minutes to lyse cells and isolate DNA. A 1500 bp region of the 16S rRNA gene was amplified with universal primers 8F (5’–AGAGTTTGATCMTGGCTCAG–3’) and 1492R (5’-CCCCTACGGTTACCTTGTTACGAC-3’). Reactions were performed in a total volume of 25 μl containing 12.5 μl 2 x MyTaq Red (Bioline, UK), 0.5 μl of each 10 μM primer stock, 1 μl template DNA, and 10.5 μl nuclease-free water. Thermal cycling conditions were 95 °C for 5 minutes, 30 cycles of 95 °C for 30 seconds, 56 °C for 45 seconds, 72 °C for 90 seconds, and a final extension at 72 °C for 7 minutes. PCR products were Sanger sequenced with forward primer 8F by GATC (GATC, Cologne) and resulting sequences were subjected to BLAST analysis (Altschul et al. 1990) against the GenBank database (http://www.ncbi.nlm.nih.gov/) to allow taxonomic identification by similarity.

### Library preparation and sequencing of the *Tatumella* sp. TA1 genome

DNA from *Tatumella* sp. TA1 was extracted as follows: Cryopreserved isolates were revived by streaking on to BHI agar and incubated at 25 °C for 72h. Single colonies were inoculated into BHI broth and incubated at 25 °C until cultures reached an OD600 of 0.3. Cultures were pelleted by centrifugation at 6,000 x g for 6 minutes. The supernatant was removed, and cells were resuspended in DNA elution buffer at a concentration of 1 x 10^5^ Colony Forming Units (CFUs) mL^-1^. DNA was extracted using the Zymo Quick DNA Universal Kit (Zymo, UK) following the manufacturer’s instructions for biological fluids and cells with the following amendments to the protocol: samples were incubated with proteinase K at 55 °C for 30 minutes rather than 10 minutes. DNA was purified with Ampure beads (Agencourt) at a 1:1 ratio, and stored at 4 °C until library preparation. DNA was sheared to 10 kb using Covaris G-tubes following the manufacturer’s guidelines, and library preparation was performed with the SMRTbell library preparation kit (Pacific Biosciences) following the manufacturer’s instructions. The Qubit dsDNA HS assay (Life Technologies, UK) was used to quantify the library, and the average fragment size was determined using the Agilent Bioanalyzer HS assay (Agilent). Size selection was performed with the Blue Pippin Prep (Sage Science) using a 0.75 % agarose cassette and the S1 marker. The final SMRT bell was purified with Agencourt AMPure XP beads (Beckman Coulter Genomics) and quantified with the Qubit dsDNA High-Sensitivity assay (Life Technologies), and an Agilent Bioanalyzer High-Sensitivity DNA chip (Agilent). The SMRTbell library was annealed to sequencing primers at values predetermined by the Binding Calculator (Pacific Biosciences), and sequencing was performed on two SMRT cells using 360-minute movie times. Pacific Biosciences sequencing and library preparation of the *Tatumella* sp. TA1 isolate was performed at the University of Liverpool Centre for Genomic Research on a Pacific Biosciences RS II sequencer.

### Genome assembly and annotation

The *Tatumella* sp. TA1 draft genome was assembled from 365,445 sub-reads with a mean length of 6,926 bp using the Hierarchical Genome Assembly Process (HGAP) workflow (Chin et al. 2013). It is available at NCBI under the accession numbers CP033727-CP033728. The HGAP pipeline comprised pre-assembly error correction of sub-reads based on read length and quality, assembly with Celera, and assembly polishing with Quiver. The assembled genome was annotated with PROKKA version 1.5.2 (Seemann 2014). To further investigate the extracellular membrane structures and mobile elements encoded in the *Enterobacter* sp. OLF, *Tatumella* sp. TA1 and three *Ca.* E. dacicola draft assemblies, genomes were re-annotated with RAST (Overbeek et al. 2014). These annotations are available in Table S3.

### Phylogenetic analyses

For the species tree, allowing phylogenetic placement of *Tatumella* sp. TA1 and Ca. *E. dacicola,* taxa were selected based on (Palmer et al. 2017), which presents a robust phylogeny of the *Erwinia, Pantoea* and *Tatumella* genera using genome-wide orthologs. Where Palmer et al. (2017) included several conspecific taxa, we chose the most biologically-relevant of the identical taxa. The Western Flower Thrip (WFT)-associated bacteria BFo1 and BFo2 were included in order to validate phylogenetic similarities with *Ca.* E. dacicola and *Tatumella* TA1 based on 16S rRNA sequences. A full list of the 46 genomes employed for this analysis can be found in Table S1A, including the 6 outgroup taxa and the 3 available *Ca.* E. dacicola draft genome assemblies (ErwSC, IL and Oroville). Genomes were annotated with PROKKA version 1.5.2 using the default settings, and 289 single-copy orthologs present in all taxa were identified using OrthoMCL version 1.4 with default parameters (Li et al. 2003). The ‘reference’ tree for the urease operon analysis was estimated from 8 genes randomly selected from the set of 289 single-copy orthologs, with a total of 5,572 informative sites; we assumed neutral evolution status for these genes based on their predicted housekeeping functions (Table S1A). For the urease subunit alpha (*ureC*) gene tree, homologs were retrieved based on BlastP queries against the nr database at NCBI (restricted to taxid=Bacteria). We included as many taxa common to the reference and species tree as possible, to ensure meaningful comparisons. Additional homologs were retrieved by targeted searches in RefSeq (full list of species names and accession numbers in Table S1B). For all trees, amino acid sequences were aligned with MUSCLE version 3.8.31 (Edgar 2004), and informative sites were selected with Gblocks version 091b (Castresana 2000). Maximum-Likelihood trees were estimated using IQ-TREE version 1.5.5 after automated model selection (Nguyen et al. 2015), with 300 random trees as burn-in. Node support was calculated using 1000 ultra-fast bootstraps (Minh et al. 2013). The multilocus coalescent-based species and reference trees were estimated from individual ortholog trees using Astral III, with node support calculated using 1000 bootstraps (Zhang et al. 2018). Trees were rooted and visualized using FigTree v.1.4.0 (http://tree.bio.ed.ac.uk/software/figtree/).

### Statistical analyses

All statistical analyses were performed in R version 3.3.3 (R Core Team 2017). Comparisons of GC content mean ranks, means and distributions were performed with Mann-Whitney, Welch’s t-test (allowing for comparison of unequal variance-samples) and Kolmogorov-Smirnov tests. For these comparisons, the GC content of the urease operon and the whole genome of *Ca.* E. dacicola (ErwSC assembly) were calculated using overlapping 50 bp windows.

### Data availability

16S rRNA amplicon sequencing data are available as part of NCBI BioProject PRJNA321174. The *Tatumella* sp. TA1 genome is available at NCBI under accession numbers CP033727-CP033728. Alignments for Figs 1 and 3 are available upon request from the corresponding authors.

## Acknowledgements

This work was supported by a BBSRC Industrial CASE Award BB/K501773/1 to ACD, awarded by the Biosciences Knowledge Transfer Network to the Food, Industrial Bioscience and Plants and crops Sectors. This work was further supported by Greek national funds through the Public Investments Program (PIP) of the General Secretariat for Research & Technology (GSRT), under the Emblematic Action “The Olive Road” (project code: 2018ΣE01300000), awarded to JV. The authors are grateful to Greg Blow and Michael Capper for reading drafts of the manuscript, and to Paschalia Kapli and Pavlos Pavlidis for discussions and technical advice on horizontal gene transfer and phylogenetic trees.

## Author Contributions

F. B., A.G., J.V. and A.C.D designed the study. F.B. and A.K. collected the samples. F.B. and I.B.G. prepared the sequencing libraries. F.B., A.G., I.B.G. M.K. and A.C.D. performed data analysis. F.B., A.G. and A.C.D. wrote the manuscript, and all authors read and made comments on the manuscript prior to submission.

## Supplementary material

**Fig. S1.** Hypothesized microbial nitrogen recycling pathways from host waste products. UO (Uricase E.C. 1.7.3.3); ALN (Allantoinase E.C. 3.5.2.5); ALC (Allantoicase E.C. 3.5.3.4); UL (Ureidoglycolate lyase E.C. 4.3.2.3); URE (Urease 3.5.1.5); GS (Glutamine synthetase E.C. 6.3.1.2).

**Table S1.** GenBank accession numbers and metadata for sequences used in phylogenetic analyses. (*A*) Whole genome sequences used for ortholog picking and phylogenetic reconstruction of the species and reference trees (B) Amino acid sequences used for the *ureC* gene tree.

**Table S2.** Presence and expression of genes related to nitrogen assimilation in *Ca.* E. dacicola and *Tatumella* sp. TA1. (*A*) Urea hydrolysis and nitrogen recycling pathways (*B*) Amino acid biosynthetic pathways.

**Table S3.** Functional annotations and corresponding sequences for the genomes of *Ca.* E. dacicola (three currently available assemblies), *Tatumella* sp. TA1 and *Enterobacter* sp. OLF, as well as for the transcriptome of *Ca.* E. dacicola (ribosomal RNAs excluded). The annotations come from RAST subsystems.

**Table S4.** Abundance of different categories of mobile genetic elements in the *Ca.* E. dacicola (three currently available assemblies), *Tatumella* sp. TA1 and *Enterobacter* sp. OLF genomes, as annotated by RAST subsystems.

**Table S5.** Functional annotation of *Ca.* E. dacicola, *Tatumella* sp. TA1 and *Enterobacter* sp. OLF genomes for extracellular surface structure components, by assignment of genes to the KEGG category “Membrane Transport” using RAST subsystems.

